# Distance to future climate analogues reveals refugia for agricultural adaptation

**DOI:** 10.1101/2025.10.31.685848

**Authors:** Anna Geldbach Jones, Samuel Pironon, Guillermo Friis, David Geldbach, James S. Borrell

## Abstract

Future climate trends are predicted to cause a dramatic shift in the geographic distribution of global agriculture. Mapping the distance to future analogues provides a crucial measure of adaptation potential, and reveals steep slope agriculture, regions with small field sizes, and ancient centres of crop domestication have more proximal climate analogues. Drawing parallels from wild biodiversity conservation, these landscapes could play a critical role as climate refugia for agrobiodiversity.

## Main

Future climate change is expected to cause poleward and altitudinal shifts in bioclimate zones^1^, generating mismatches between crop distributions and climates suitable for their production^2^. This threatens global yields with evidence of slowed agricultural productivity growth already emerging^3,4^. However, exposure to climate change varies greatly across the Earth’s >600 million farms^5,6^. We suggest that where climatic change is predicted, persistence and adaptation may be more likely if an analogous climate-zone is nearby in the future. This proximity can be measured in terms of the ‘climate distance’.

We calculate the distance to an area’s closest future climate analogue, climate distance, in terms of annual temperature and rainfall for 10 million randomly sampled 1 km^2^ cells (approximately 2% of global land), between 2011-2040 and 2040-2070, under the SSP370 scenario, projected with GFDL-ESM413 (Fig 1a). Climate distance is proportional to climate velocity^7^, the rate species must migrate at to remain within a climate niche, but for agriculture we suggest the impacts of climate shifts are described better by distance than velocity, since adaptation through adoption of alternative crops is mediated by proximity rather than migration rates. In the resulting global map, we find large areas of the highly productive Russian/Asian steppe have high climate distances, illustrating that to maintain productivity, alternative crops, agronomic practices, and expertise would have to be sourced from much further away (Fig 1a). Conversely large areas of Europe, Western North America, and Northern Africa have shorter climate distances, enabling adoption of agricultural practices from more proximal, neighbouring regions.

**Figure 1.**
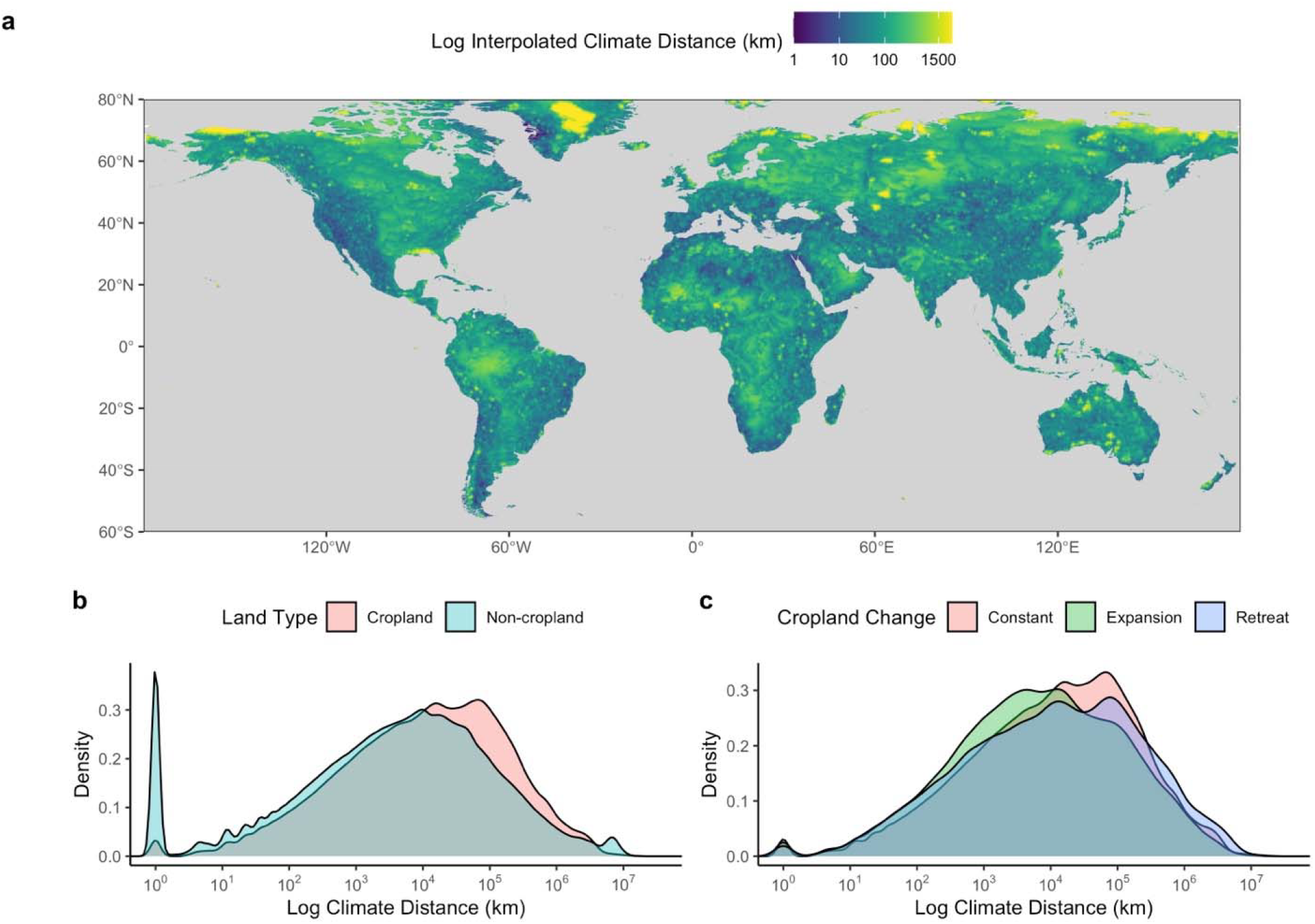
Global patterns of climate distance, illustrating a) regional patterns, b) differences between cropland and non-cropland, and c) changes in cropland climate distance associated with the expansion and contraction of agricultural land use.

We expect climate distance to be strongly mediated by topography and slope (Supplementary Material 1). Sloped areas have steeper spatial gradients of climate, meaning future climate analogues will be found close by along the gradient^7^. Wang et al^8^ reported that future climate shifts disproportionately threaten steep-slope agriculture using Koppen–Geiger classifications at 1 km resolution for the present (1980-2016) and future (2071-2100). The authors concluded that steep-slope agriculture is more likely to change climate classification than average global agricultural lands and is therefore at risk. However, our data suggests important nuance to this interpretation, illustrating that agricultural land on steep slopes has significantly shorter distance to climate analogues than flat agricultural lands (Fig 2a, steep cropland mean = 56 km, non-steep cropland mean = 103 km, Wilcoxon rank-sum test W = 1.79 × 10□, *P* < 0.001). This is an important mediator of resilience, because the impact of bioclimatic shifts on food systems depends on farmers’ ability and capacity to either move with shifting climate or adapt *in situ* by adopting new agricultural practices. Therefore as agriculture in hilly and mountainous areas will likely be subject to more proximal climate shifts, this will potentially enable farmers more time to adapt, and require adoption of crops or practices from closer regions.

**Figure 2.**
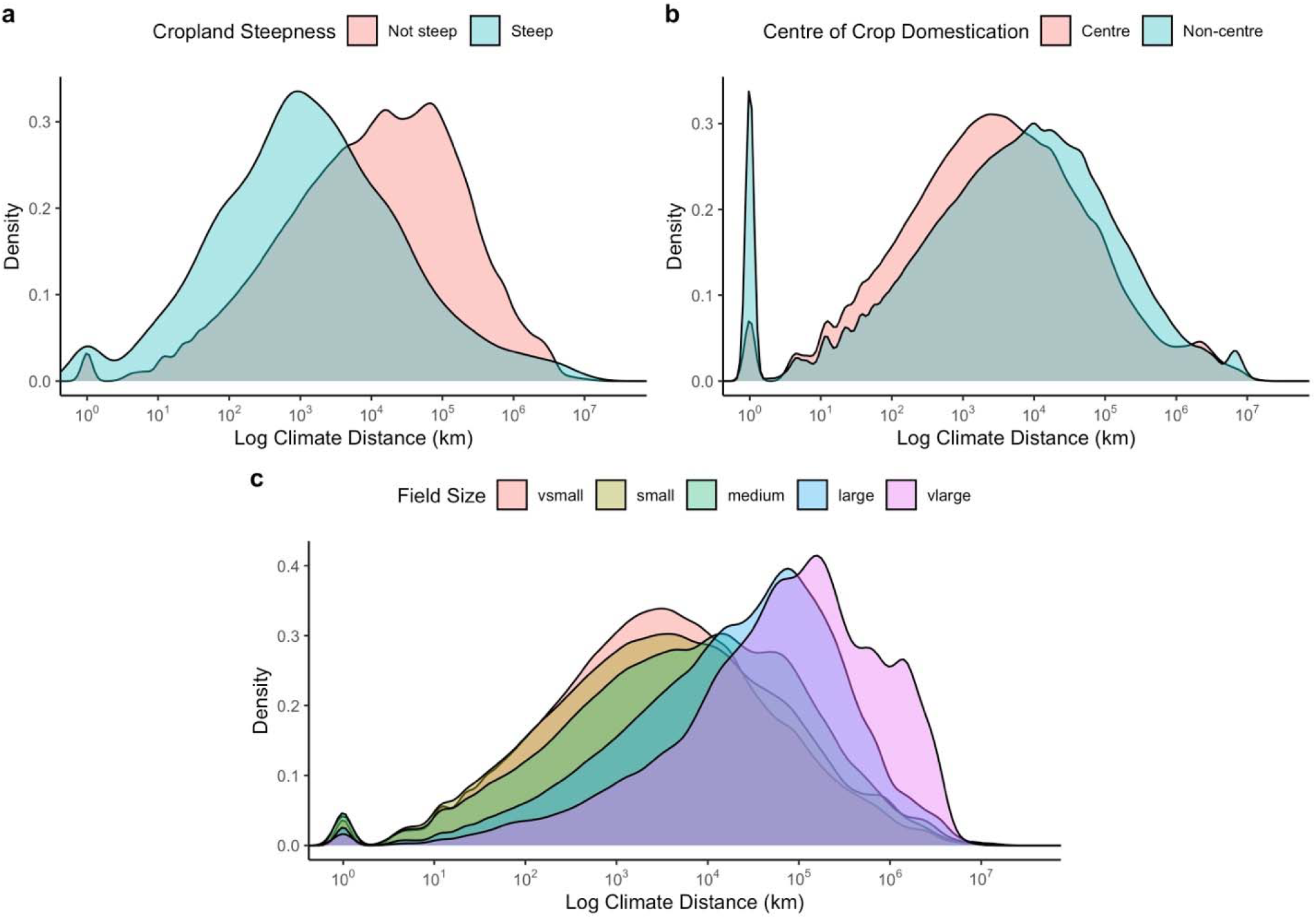
Climate distance across a) steep versus not steep cropland, b) centres and non-centres of domestication and c) agricultural field size.

Previous analyses identified that rates of adoption by farmers of climate-resilient crops and varieties has been highly variable^10,11^. Important socioeconomic determinants include effective agricultural extension services, education level, and crucially geographic factors such as farmers’ access to inputs like seeds and relevant information^10,11^. To compare proximity to climate analogues across a gradient of commercial to smallholder farming systems, we examined how climate distance varied with dominant field size^12^ and found a significant association (χ^2^ = 159232, df = 4, *P* < 0.001). Post hoc testing showed climate distance increased significantly with larger field size (Dunn’s test with Benjamini-Hochberg correction, all *P* < 0.001). We also found that the proportion of 1 km^2^ areas where the climate did not change between time points decreased with increasing field size (Cochran-Armitage test for trend, Z = 13.77, dim = 5, *P* < 0.001). Therefore, smaller farms are less likely to experience changing climate, but if they do, future climate analogues and therefore potential new crops will be nearer.

Conversely areas with larger field sizes, often associated with commercial agricultural systems, had greater climate distances. This has important food system implications, because although small and medium farms produce the majority of food globally, certain regions including North America, Australia, and New Zealand depend on large farms for over three quarters of their production^13^.

Reflecting on these pattners, we hypothesise that climate distance has helped shape the global distribution of agriculture. Consistent with this, we find the proportion of 1 km^2^ areas where climate is not predicted to change between time points (climate distance = 0) was nearly ten times less in cropland compared to non-cropland (cropland = 0.6%, non-cropland = 5.7%, χ^2^ = 45113, df = 1, *P* < 0.001) illustrating the stability of these areas. However, where the climate *is* predicted to change, cropland is further from future climate analogues than non-cropland (Fig 1b, cropland mean = 103 km, non-cropland mean = 92 km; Wilcoxon rank-sum W = 4.01 × 10^12^, *P* < 0.001).

We also examined whether recent changes in the distribution of croplands are projected to alter exposure to climate change. The global cropland expansion rate has nearly doubled in the last two decades^18^, occupying around 12% of land in 2021^19^. We compared the climate distance calculated in our analysis for areas where cropland has expanded, stayed constant, or retreated between 2003-2019 and found significant differences^20^ (Fig 1c, Kruskal-Wallis, χ^2^ = 3801.5, df = 2, *P* < 0.001). New areas converted to cropland have shorter predicted climate distances than constant or retreat areas (expansion mean = 89 km, constant mean = 104 km, retreat mean = 113 km; post hoc Dunn’s test expansion versus constant Z = 61.64, *P* < 0.001, expansion versus retreat Z = -33.64, *P* < 0.001). Similarly, there were significantly more areas where climate was not predicted to change in regions of cropland expansion compared to constant cropland regions (pairwise comparison, *P* = 0.0027, Bonferroni-adjusted). This suggests new areas of cropland will be less exposed to climate change and better able to adapt via more proximal climate analogues in the future, potentially providing early evidence of food system adaptation.

As climate change accelerates in coming decades, climate refugia – areas that remain relatively buffered from climate change – will become increasingly important for species persistence^5^. Incorporating refugia into spatial conservation planning is one tool for mitigating global biodiversity loss^14^. This applies to both wild and agrobiodiversity, as well as bio-cultural knowledge^15,16^. Areas with more proximal climate analogues will enable a slower rate of crop turnover, more time for adaptation, and transmission of cultivation practices from closer sources. Ancient centres of crop domestication are regions where most of the world’s major crops originated, and today contain globally important agrobiodiversity which underpins the resilience of our food systems. We find significantly shorter average climate distance within centres of domestication than land outside centres of domestication (centres of domestication mean = 72 km, non-centres = 93 km, Wilcoxon test W = 4.88 × 10^12^, *P* < 0.001). Where climate change is projected to disrupt agricultural systems, our results suggest that centres of domestication may be suitable targets for conservation as refugia. Considering long-term climate stability is also often associated with high endemism^17^, the shorter predicted climate distance of centres of crop domestication may mirror historical stability underpinning the development of novel crops.

To test sensitivity and uncertainty in our modelling choices we performed three analyses. First, we examined the importance of an appropriate climate threshold by mathematically simulating climate distance (Supplementary Material 1). Threshold refers to the maximum change in climate for an area to still be classed as climatically analogous. If the threshold is too small (higher climatic resolution), future climate analogues are far away due to discretisation and steep areas are disproportionally affected by climate input noise. Conversely if the threshold is too large, nuanced climate variability is lost and fewer climate shifts are detected. An appropriately chosen threshold will be greater than the magnitude of climate prediction noise, and smaller than true climate variability. Practically, we show that this can be approximated as an order of magnitude less than the standard deviation of the climatic variables. Second, we considered the effect of using different climate models as inputs (Supplementary Material 2). We demonstrate highly significant rank order agreement between models (W=0.999, χ2(8591) = 34327, p<0.001 for temperature and W = 0.997, χ2(8591) = 34262, p<0.001 for precipitation) indicating that relative climate distance has low sensitivity to model choice (Supplementary Material 2).

Many of the world’s “breadbasket” regions are particularly exposed to climate change due to long distances to future climate analogues. This illustrates the growing need for transfer of crops and knowledge across landscapes to alleviate production losses^9^. Traditional, smallholder dominated agricultural landscapes buffered from climate change by their topography are promising candidate refugia for agrobiodiversity, and sources of replacement crops or varieties^22^. A strategy of area-based agrobiodiversity conservation targeted towards climate refugia in ancient centers of crop domestication, and underpinned by appropriate support, is needed to safeguard these sites and improve the climate resilience of global agriculture.

## Methods

To estimate climate distance globally, we collated annual mean annual air temperature and annual precipitation data from CHELSA BIOCLIM at 1 km resolution for T1 (2011-2040) and T2 (2040-2070) under SSP3-RCP7.0 (SSP370) scenario, projected with GFDL-ESM4^23^. GFDL-ESM4 was used because it has highest priority in the ISIMIP3b protocol, a comparison with other global climate models is presented in Supplementary Material 2. We randomly sampled 2% of global terrestrial 1 km cells (n = 10,000,000), and for each cell calculated the minimum Haversine distance to a cell in T2 which had analogous climate variables to the focal cell in T1, up to a maximum distance of +-10 degrees in latitude and longitude. Based on the global standard deviation of annual temperature and rainfall in T2 (temperature SD = 10.5, rainfall SD = 843), we chose a threshold of ±0.5 °C annual temperature change and ±5 mm annual rainfall change to define cells which experienced a climate shift between T1 and T1. The annual temperature and rainfall data was rounded to zero decimal places, and we then performed exact matching for temperature and matched rainfall to ±5 mm. For further discussion of the appropriate climatic thresholds for investigating climate distance based on simulations and well as mathematical characterisation of the effects of slope on climate distance, see supplementary material.

To make a global map of climate distances, we used linear interpolation to create a continuous visualization based on a random 1 million climate distance results. For visualisation as a global map, where no climate analogue was found, the climate distance was set to 3000 km. A log+1 transformation was applied for visualization. Based on our climate distance results for a 2% sample, we also predicted climate distance at 1 km resolution for the entire world using a random forest algorithm which used temperature, rainfall, slope, latitude and longitude as predictors. The random forest prediction algorithm achieved an R^2^ of 0.94 when tested on the withheld 20% of our calculated climate distance data. This global map of predicted climate distance at 1 km resolution is the first of its kind to be publicly available and could be used to answer many other pertinent questions about climate change exposure beyond agriculture, for example in reforestation, conservation or future proofing urban biodiversity. In our own analysis of agricultural land types, we used calculated climate distance based on 2% of the world rather than predicted data.

Cropland extent timeseries for the period 2003-2019 were derived from Potapov et al, and down sampled from 300 m to 1 km resolution^18^. We calculated slope from the GTOPO30 digital elevation map^24^ and used a slope of 12% (∼6.8 °) to define steep slope agriculture, consistent with previous analyses^8^. Global field size estimates were derived from Lesiv *et al*.^12^ who used crowd sourced visual interpretation of high resolution satellite imagery. Known centres of crop domestication were adapted from Maxted and Vincent^25^. Climate distance data was right skewed with many zero values, so non-parametric tests were used and log+1 transformations applied to visualisations. We used Morran’s I test to check for spatial autocorrelation and then applied Wilcoxon signed-rank test to examine differences in density distributions of climate distance between groups. Where three groups were compared we used the non-parametric Kruskal-Wallis test followed by Dunn’s test using Bonferroni-adjusted *P*-value for multiple pairwise comparison. All analyses were conducted using R (version 4.3.1) or Python (version 3.11).

## Data and Code Availability

The code for calculation of climate distance used, as well as subsequent analyses are available on Github. The calculated climate distance dataset is available on Zenodo. The predicted 1km global map of climate distance is available via a public <u>Google Earth Engine script</u> and as an <u>asset</u>.

