## Supplementary Materials 1 for "Distance to future climate analogues reveals refugia for agricultural adaptation"

### The effect of steepness on climate distance

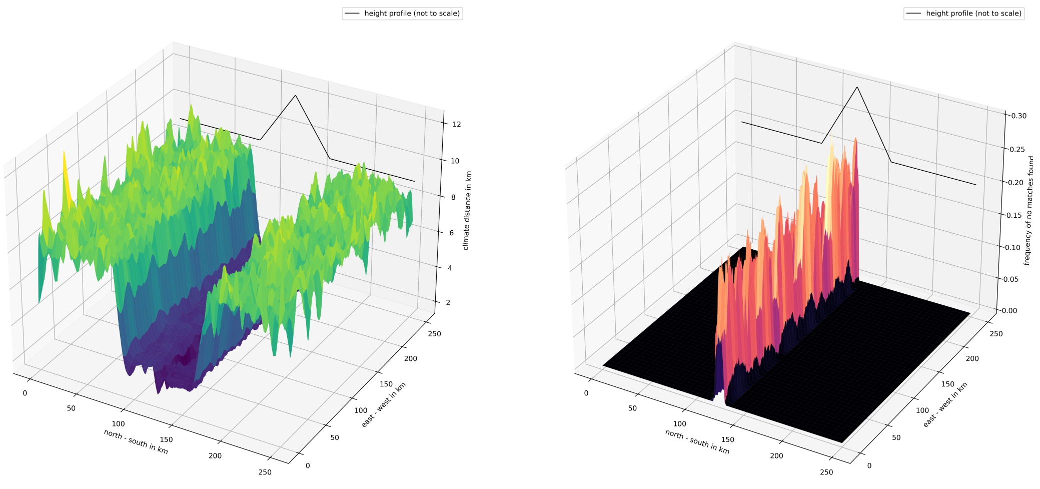


Figure 1: Simulation of climate distance in the presence of a mountain range with a maximal altitude $1500m$. The climate distances are smaller on the slopes than on the plains but we do not find future climate analogues for the top of the mountain range within the simulation range.

##### Simulating climate distance

We would like to understand the effect of steep areas (hills or mountains) on climate distance using simulated data. We do this by simulating temperature values before and after climate change. Suppose we have a $N \times N$ grid $\Lambda$ and a function $h : \Lambda \to [0, 1]$ that measures the relative altitude of a given point on the grid. We model the temperature of a lattice point $(x, y) \in \Lambda$ pre-climate change as

$$T_{old}(x, y) = t_{base} - \alpha \frac{x}{N} - \beta h(x, y) + \varepsilon_{env}(x, y), \tag{1}$$

where $t_{base}, \alpha, \beta \geq 0$ are parameters and where $\varepsilon(x, y)$ is random noise.

1. $t_{base}$ is a reference temperature.
2. $\alpha$ models the north–south gradient in temperature.
3. $\beta$ models the dependency of temperature on altitude, equivalently $\beta$ models the maximal altitude of the mountain range.
4. $\varepsilon(x, y)_{env}$ is environmental noise due to climate variability. We assume that $\{\varepsilon_{env}(x, y)\}_{(x,y)\in\Lambda}$ are independent random uniform random variables on $[0, 2\mu_{env}]$.

After climate change, the new temperature is given by

$$T_{new}(x, y) = T_{old} + \delta + \varepsilon_{pred}(x, y), \tag{2}$$

where $\delta \geq 0$ and $\varepsilon_{pred}(x, y)$ is random noise.

5. $\delta$ models the strength of change in temperatures due to climate change.
6. $\varepsilon(x, y)_{pred}$ capture both uncertainty of climate modelling and measurement errors. We assume that $\{\varepsilon_{env}(x, y)\}_{(x,y)\in\Lambda}$ are independent uniform random variables on $[0, 2\mu_{pred}]$.

A reasonable choice of altitude function $h$ is given by

$$h(x,y) = \max\left\{1 - \frac{|x/N - 1/2|}{10}, 0\right\}, \tag{3}$$

which models a mountain range in east–west direction. We can run simulations like this but they are severely limited in the scaling parameter $N$ as the algorithm to determine climate distance has a run time of $O(N^4)$.

In this special case of simulated data we have access to hierarchical search methods that allow for a quicker computation of climate distances.[1] We plot the results in Figure 1. We discuss the choice of parameters (6) in the following sections.

##### Reducing it to a one–dimensional model

To improve the performance of the simulations, we make use of the east–west symmetry of this model. If we look for a climate analogue of a location $(x, y)$, then the analogue will likely have a very similar $y$–coordinate even if it has a very different $x$–coordinate. We therefore restrict ourselves to a one–dimensional model where $\Lambda = \{1, \ldots, N\}$ models the north–south direction. This reduces the run time of the simulation to $O(N^2)$ which greatly increases out ability to run simulations. We use

$$h(x) = \max\left\{1 - \frac{|x/N - 1/2|}{10}, 0\right\}, \tag{4}$$

to model the shape of the cross–section of the mountain range. When searching for climate analogues, we call $(x', y')$ a climate analogue for $(x, y)$ if

$$|T_{new}(x', y') - T_{old}(x, y)| \leq \gamma, \tag{5}$$

where the threshold $\gamma > 0$ is a parameter.

##### Steep vs flat areas in simulation

We simulate this with the following parameters:

$$t_{base} = 30, \quad \alpha = 10, \quad \beta = 10, \quad \delta = 1, \quad \mu_{env} = 1, \quad \mu_{pred} = 0.1, \tag{6}$$

and two different choices for the matching threshold $\gamma$,

$$\gamma \in \{0.2, 0.02\}. \tag{7}$$

Using that the north–south temperature gradient is approximately $1/150\ C/km$ in temperate regions and the altitude gradient is approximately $1/150\ C/m$, this corresponds to mountain range with maximal altitude $1500m$ and a width of $300km$. In Figure 2 we simulate the climate distance.

We observe several things here.

1. If the threshold is chosen properly, areas of high slope have a lower climate distance than flat areas.
2. As expected, we do not find a climate analogue for the top of the mountain.
3. If we choose the threshold $\gamma$ too small, there are more locations where we do not find a future temperature match. Here the high–slope areas are disproportionately affected.
4. Similarly, due to the random noise overpowering the threshold, locations with high slope now do not find matches close by. Rather, they are matched with locations very far away in the flat areas leading to high climate distance.

This means that choosing a threshold that is too small falsely reverses the effect of slope on climate distance.

[1] Theoretically, this could achieve a computational complexity of $O(N^2 \log(N))$, practically it is closer to $O(N^3)$. Nevertheless, this is a significant improvement.

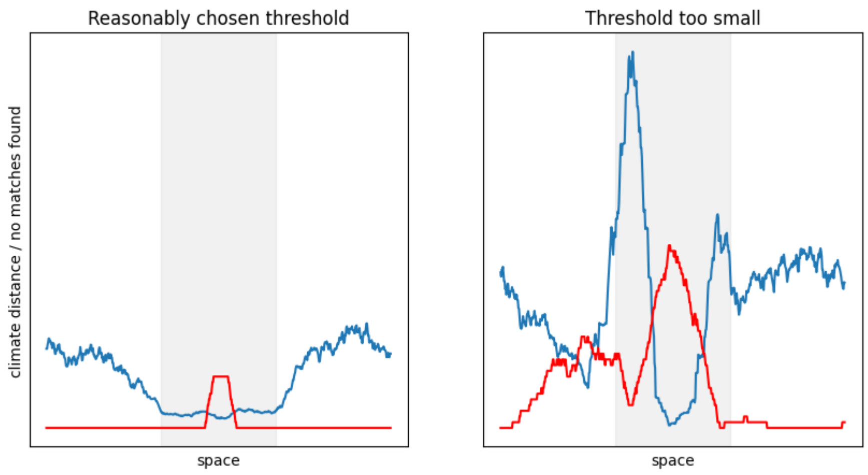


Figure 2: Simulating climate distance for a mountain range with two different thresholds for when two temperatures count as match. The mountain range is shaded in grey. The climate distance is plotted in blue, in the local frequency of points with no match found in plotted in red.

##### Choosing the right threshold

It is also interesting to look at mean climate distance as a function of the threshold, see Figure 3. Here, if for a steep location $x$ we did not find a future climate analogue, we assigned the maximal observed distance over steep locations to its climate distance,

$$\text{climate distance}(x) = \max\{\text{climate distance}(y) : y \text{ is steep and we found an analogue for } y\}. \tag{8}$$

This captures the effect that for small thresholds there are plenty of locations where we do not find an analogue. Nevertheless, we do not want to assign an arbitrary value or skip these locations. This simulation again shows that the threshold needs to be chosen carefully and not be too small.

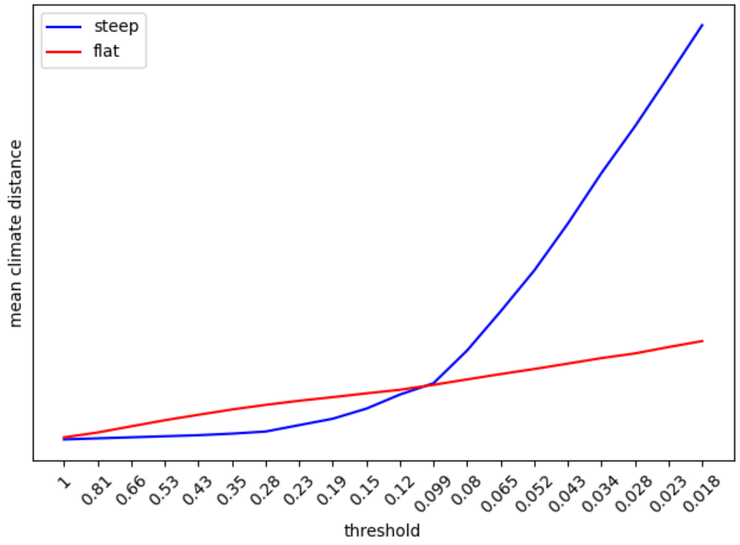


Figure 3: Simulated mean climate distance as as a function of the threshold. A lower threshold corresponds to a higher resolution in the search algorithm.

##### Details on the threshold

The performance of the climate distance algorithm depends strongly on the choice of threshold $\gamma$ in (5). This is related to the discretisation of space. Suppose that our discrete space $\Lambda$ approximates a continuous space $[0,1]$ and that $T_{new}$ and $T_{old}$.

In this framework, the climate distance algorithm corresponds to finding $y \in [0,1]$ such that $T_{new}(y) = T_{old}(x)$ for a given $x \in [0,1]$. We introduce the threshold $\gamma$ to model that we are not looking for an exact match, but rather an approximate match up to an error $\gamma$. In Figure 4, this means that we are looking for points $y$ where $T_{new}(y)$ lies in the grey strip. In this example, there are many approximate matches as $f_{new}$ crosses the strip multiple times.

The issue now is that we are not working with $[0,1]$ but with a discretised space. This means that we need to choose $\gamma$ big enough so that it is unlikely that a match will be missed because the discretisation is too coarse. In Figure 4 this happens as none of the lattice points fall into the grey strip for the two later crossing. This could happen in a situation where for $x, y \in \Lambda$

$$\begin{cases} T_{new}(y) + \gamma < T_{old}(x) < T_{new}(y+1) - \gamma, \\ |\varepsilon_{env}(y) - \varepsilon_{env}(y+1)| \quad \text{is small.} \end{cases} \tag{9}$$

Here neither $y$ nor $y+1$ are a match for $x$ but because we would assume that $T_{new}$ is continuous (because $|\varepsilon_{env}(y) - \varepsilon_{env}(y+1)|$, the local climate variability, is small) there should be a match somewhere between $y$ and $y+1$. This match is missed by due to the discretisation.

In the context of (1) and (2), we want to choose $\gamma$ so that with high probability

$$\varepsilon_{pred} < \gamma < \varepsilon_{env}. \tag{10}$$

This ensures that no matches are lost to the noise of climate prediction while at the same time we do not lose the effect of the noise from climate variability. We have chosen $\varepsilon_{pred}$ and $\varepsilon_{env}$ so that their expected values are $\mu_{pred}$ and $\mu_{env}$. This means that a reasonable restriction on $\gamma$ is

$$\mu_{pred} < \gamma < \mu_{env}. \tag{11}$$

Because we used uniform random variables, $\mu_{pred} = c\sigma_{pred}$ and $\mu_{env} = c\sigma_{env}$, where $\sigma_{pred}, \sigma_{env}$ are the standard deviations and where $c \approx 0.58$ is a universal constant. In the context of the simulations of Figure 2 and 3, this means $0.1 < \gamma < 1$. In Figure 2, the first simulation used $\gamma = 0.2$ and the second simulation uses $\gamma = 0.02$, see (7). This is aligned with (11) which suggests that $\gamma = 0.2$ is a good choice. Lastly, we want to remark that because $\varepsilon_{env}$ has a higher variance than $\varepsilon_{pred}$, we want $\gamma$ to be towards the lower end in the interval $\gamma \in [\mu_{pred}, \mu_{env}]$.

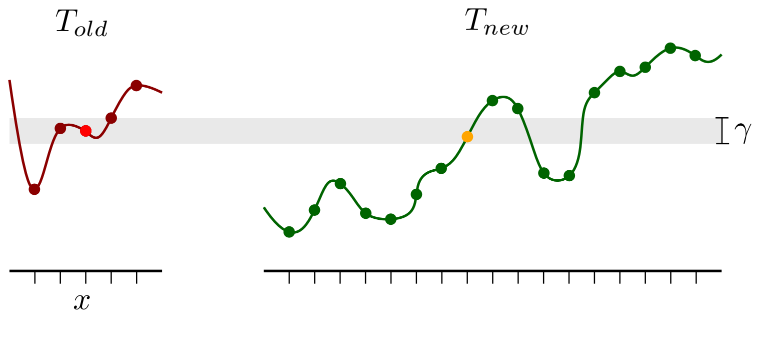


Figure 4: An illustration of the threshold. We find one match for $x$ which is the point in orange. Note that there are more potential matches that are narrowly missed because $\gamma$ is small.
