## Supplementary Materials 2 for "Distance to future climate analogues reveals refugia for agricultural adaptation"

### Supplementary Material 2

CHELSA V.2.1 is available for a range of CMIP6 scenarios and climate models. The global climate models are selected based on the intersectoral impact model intercomparison project (ISIMIP) 3b, see CHELSA V2.1 Technical Specification for further details. The GFDL-ESM4 model data was used in our main analysis, which has the highest priority in the ISIMIP3b protocol.

To better understand the role of climate model choice as a source of uncertainty in our climate distance calculations, we examined the correlations between GFDL-ESM4 and three alternative models under SSP370: IPSL-CM6A-LR, MPI-ESM1-2-HR and UKESM1-0-LL.

We first present a global comparison of climate models, masked to land area. Then additionally, to provide a more meaningful comparison, and avoid over estimating correlations as a result of including global temperature extremes, we restricted our evaluation to a region of East Africa (~10 degrees x ~20 degrees) containing a wide range of topography similar to the scale of our main analysis. Using randomly sampled 1km cells, the four models showed very close agreement for average temperature and annual rainfall at both global (Figure 1) and regional (Figure 2) scales.

In our analysis, the most important measure of correlation is rank order, because while the overall magnitude of climate change may differ between models and depending on the future time scale studied, rank order of cells determines whether the same regions have lower or higher distance to future climate analogy. Therefore we calculate global pairwise Spearman’s rank correlation for temperature (Table 1) and precipitation (Table 2) respectively. We also estimate global Kendall’s W to test for rank-order concordance between multiple models.

Spearman’s Rank Correlation between the models revealed correlations consistently >0.99 for both temperature and precipitation. Kendall’s coefficient also indicated **very high agreement**, W=0.999, χ2(8591) = 34327, p<0.001 for temperature and W = 0.997, χ2(8591) = 34262, p<0.001 for precipitation. Therefore we are confident that our conclusions are robust to alternative future climate projections.

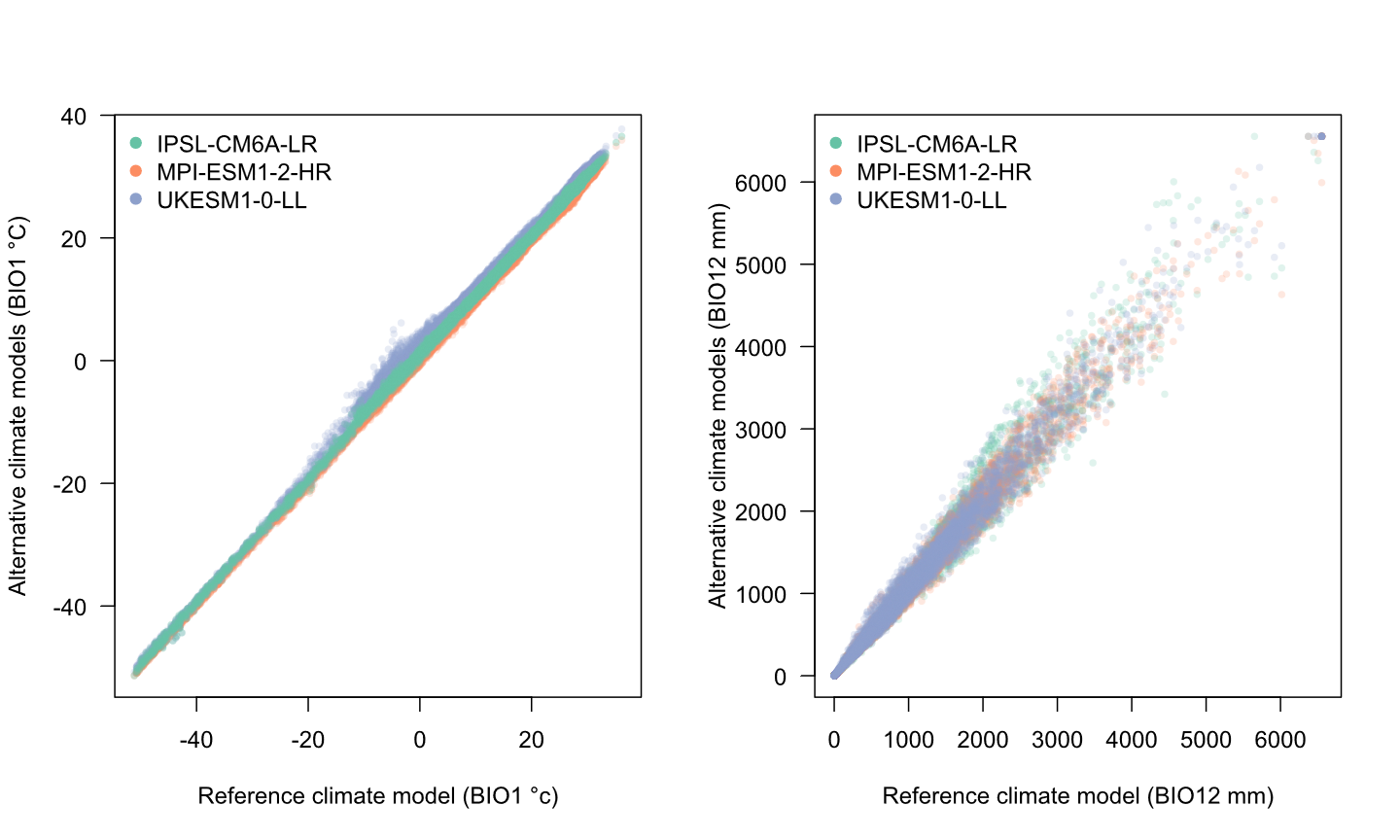

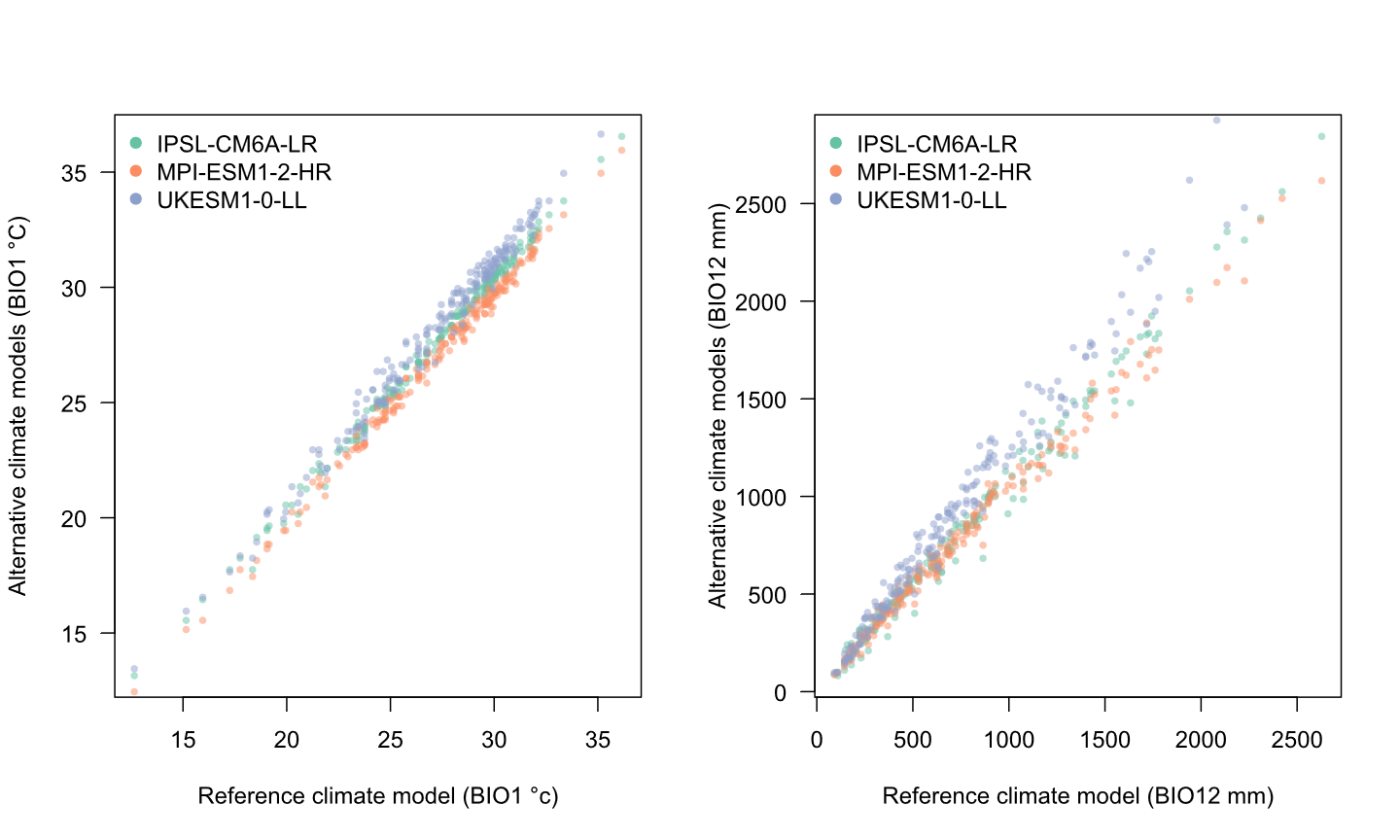
**Figure 1:** Correlation of different global climate model predictions of annual average temperature (°C, BIO1) and annual rainfall (mm, BIO12) to the GFDL-ESM4 predictions used in our analyses for 10,000 randomly sampled 1km cells.

**Figure 2:** Correlation of different global climate model predictions of annual average temperature (°C, BIO1) and annual rainfall (mm, BIO12) to the GFDL-ESM4 predictions used in our analyses for 200 randomly sampled 1km cells within a the region defined by a bounding box (29.5, 48.0, -5.0, 15.0).

**Table 1.** Spearman’s rank correlation across four climate models for annual mean temperature.

| Global analysis - BIO1 | gfdl-esm4_ssp370 | ipsl-cm6a-lr_ssp370 | mpi-esm1-2-hr_ssp370 | ukesm1-0-ll_ssp370 |
| --- | --- | --- | --- | --- |
| gfdl-esm4_ssp370 | 1.000 | 0.999 | 0.999 | 0.999 |
| ipsl-cm6a-lr_ssp370 | 0.999 | 1.000 | 0.999 | 0.998 |
| mpi-esm1-2-hr_ssp370 | 0.999 | 0.999 | 1.000 | 0.998 |
| ukesm1-0-ll_ssp370 | 0.999 | 0.998 | 0.998 | 1.000 |

**Table 2.** Spearman’s rank correlation across four climate models for annual precipitation.

| Global analysis - BIO12 | gfdl-esm4_ssp370 | ipsl-cm6a-lr_ssp370 | mpi-esm1-2-hr_ssp370 | ukesm1-0-ll_ssp370 |
| --- | --- | --- | --- | --- |
| gfdl-esm4_ssp370 | 1.000 | 0.997 | 0.997 | 0.995 |
| ipsl-cm6a-lr_ssp370 | 0.997 | 1.000 | 0.998 | 0.996 |
| mpi-esm1-2-hr_ssp370 | 0.997 | 0.998 | 1.000 | 0.995 |
| ukesm1-0-ll_ssp370 | 0.995 | 0.996 | 0.995 | 1.000 |
